# Therapeutic targeting of SLC6A8 creatine transporter inhibits KRAS mutant and wildtype colon cancer and modulates human creatine levels

**DOI:** 10.1101/2021.04.26.441371

**Authors:** Isabel Kurth, Norihiro Yamaguchi, Celia Andreu-Agullo, Helen S. Tian, Subhasree Sridhar, Shugaku Takeda, Foster C. Gonsalves, Jia Min Loo, Afsar Barlas, Katia Manova-Todorova, Robert Busby, Johanna C. Bendell, James Strauss, Marwan Fakih, Autumn J. McRee, Andrew E. Hendifar, Lee S. Rosen, Andrea Cercek, Robert Wasserman, Scott L. Spector, Syed Raza, Masoud F. Tavazoie, Sohail F. Tavazoie

## Abstract

Colorectal cancer (CRC) is a leading cause of cancer mortality. Creatine metabolism was previously shown to critically regulate colon cancer progression. We report that RGX-202, an oral small-molecule SLC6A8 creatine transporter inhibitor, robustly inhibits creatine import in vitro and in vivo, reduces intracellular phosphocreatine and ATP levels and induces tumor cell apoptosis in CRC. RGX-202 suppressed tumor growth across KRAS wild-type and KRAS mutant xenograft, syngeneic and patient-derived xenograft colorectal cancers. Anti-tumor efficacy correlated with tumoral expression of creatine kinase B. Combining RGX-202 with 5- fluorouracil or the DHODH inhibitor leflunomide caused regressions of multiple colorectal xenograft and PDX tumors of distinct mutational backgrounds. RGX-202 also perturbed creatine metabolism in metastatic CRC patients enrolled in a Phase-1 trial, mirroring pharmacodynamic effects on creatine metabolism observed in mice. This is, to our knowledge, the first demonstration of pre-clinical and human pharmacodynamic activity for creatine metabolism targeting in oncology, revealing a critical target for CRC.

## Introduction

Colorectal cancer is the second leading cause of cancer death in men and women in the United States (*1*) and a major cause of mortality worldwide. In 2020, roughly 148,000 individuals were diagnosed with colorectal cancer and 53,200 died from the disease. Early-stage colorectal cancers are primarily treated by surgical resection. For larger tumors and those that have spread to lymph nodes, 5- fluorouracil based chemotherapy regimens are administered in the post-surgical ‘adjuvant’ setting to reduce the risk of metastatic relapse. For cancers that have spread to distant organs such as the liver— the primary site of metastatic relapse—systemic 5-fluorouracil containing chemotherapy regimens are given. Targeted therapies including antibodies that engage the Epidermal Growth Factor Receptor (EGFR) or Vascular Endothelial Growth Factor (VEGF) can in combination with chemotherapeutics provide modest survival benefits (*2, 3*). However, most metastatic patients succumb to their disease, with a meager 5-year survival of 14% (*1*). Recently, innovative small molecules that covalently bind and inhibit the KRAS G12C oncogenic driver variant have elicited clinical regression responses in patients harboring this mutant allele (*4–7*). Other approaches include the development of inhibitors targeting down-stream KRAS signaling in RAS-mutant tumors (*8, 9*). Limitations of such therapies include the emergence of resistance (*10*), their lack of efficacy against other KRAS mutant alleles present in ∼40% of CRC patients, and the expected lack of efficacy in KRAS wildtype tumors (*11*). Additional approaches are thus needed to tackle the breadth of colorectal cancer.

Cancers evolve multiple mechanisms to sustain proliferative growth and survival. One such adaptive mechanism is altered metabolism, commonly referred to as metabolic rewiring. By altering the flux of metabolites in various metabolic pathways, cancer cells enhance biosynthesis of anabolic building blocks required for growth such as nucleotides, amino acids and lipids (*12*). Certain metabolites, however, can become limiting during cancer progression—requiring their extracellular import through metabolic transporters. The search for such metabolic dependencies is an active area of cancer research (*13, 14*).

We previously reported that during progression of colorectal cancer to liver metastasis, cancer cells upregulate creatine kinase brain-type (CKB) and secrete CKB into the extracellular space (*15*). CKB phosphorylates the metabolite creatine using ATP, thereby generating phosphocreatine. The γ- phosphate of phosphocreatine has ∼50% greater free energy than the γ-phosphate of ATP (*16*). Phosphocreatine thus serves as a rapidly mobilizable high-energy phosphate reserve that can yield ATP in a reaction that does not require oxygen. Because of its high energy content, phosphocreatine is stored at high levels in metabolically active tissues such as muscle, brain and kidney—allowing tight maintenance of organismal ATP levels, which are critical for a vast number of metabolic and homeostatic processes (*17*). Although extracellular ATP levels are normally low, the death of cancer cells and stromal cell substantially increases extracellular ATP concentrations in the tumor microenvironment, which have been observed to exceed 700 μM (*18*). Excess of ATP over circulating creatine levels, which range from 9-90 μM, thus favors phosphocreatine generation in the tumor microenvironment—a reaction mediated by tumor-secreted CKB. Phosphocreatine import by SLC6A8 enhanced tumoral ATP levels and accordingly promoted cell survival under hypoxia (*15*). Importantly, extracellular phosphocreatine supplementation rescued the metastatic defect and *in vitro* survival phenotype of CKB-depleted cells (*15*). Gastrointestinal tumors, such as CRC and pancreatic cancers are highly hypoxic (*19–21*), as are metastases formed by these cancers. This metabolic axis provides a mechanism to support tumor growth in hypoxic environments. Importantly, the expression levels of CKB and SLC6A8 are associated with increased colorectal cancer metastasis in patients (*15*).

Here, we identify the SLC6A8 transporter as a therapeutic target in colorectal cancer. We show that a small-molecule creatine mimetic (β-guanidinoproprionic acid, also known as RGX-202) significantly inhibits SLC6A8 and suppresses the growth of colorectal cancer tumors in syngeneic, xenograft and PDX models and caused tumor apoptosis *in vivo*. RGX-202 also strongly suppressed liver metastasis formation. Importantly, we show that this therapy is effective in both KRAS wildtype tumors and tumors bearing various KRAS mutations, including the KRAS G12D allele, which is not currently druggable by clinical stage KRAS inhibitors. Anti-tumor efficacy by RGX-202 correlated with increased CKB expression in tumors. RGX-202 exhibited greater activity in combination with multiple standard of care regimens, including 5-FU and gemcitabine, relative to single agent efficacy. We also show that RGX-202 synergizes with the dihydroorotate dehydrogenase (DHODH) enzyme inhibitor leflunomide, an oral compound previously shown to suppress colorectal cancer cell growth under hypoxia by repressing nucleotide biosynthesis (*22*). Metabolic profiling studies revealed that RGX-202 suppresses intracellular phosphocreatine, creatine and ATP levels. Finally, we observe a drug exposure-dependent increase of creatine in the blood and urine of mice and in patients treated with oral RGX-202 therapy, confirming creatine transporter inhibition in patients and supporting further clinical testing of RGX-202 in later stage clinical trials.

## Results

### Oral RGX-202 suppresses *in vivo* creatine import and depletes cellular phosphocreatine and ATP levels

β-Guanidinoproprionic acid (β-GPA, RGX-202) is a creatine mimetic that competitively inhibits the creatine transporter SLC6A8 (*23, 24*). We have developed a compressible salt-form of β- guanidinoproprionic acid (RGX-202-01) suitable for oral administration. To assess the degree to which RGX-202 inhibits SLC6A8 *in vivo*, we monitored creatine import into cardiac tissue, which expresses the SLC6A8 transporter for creatine/phosphocreatine import and energetic buffering (*17, 25*). We administered increasing concentrations of RGX-202, followed by injection of deuterium labeled creatine (d3-creatine) into wild-type and SLC6A8 knock-out mice. Heart tissue was extracted and analyzed for levels of d3-creatine by LC-Mass Spectroscopy (LC-MS/MS) (*23*). RGX-202 treatment inhibited uptake of cardiac d3-creatine in a dose-dependent manner by up to 75% at 500 mg/kg (Fig. 1a). D3-creatine levels in SLC6A8 knock-out animals were below the lower limit of quantification, confirming that creatine import is exclusively mediated by SLC6A8 (Fig. 1a). To determine if RGX- 202 could inhibit tumoral SLC6A8, we conducted similar studies in mice bearing syngeneic UN-KPC- 961 pancreatic tumors. RGX-202 at 800 mg/kg supplemented in the diet for 35 days indeed suppressed tumoral d3-creatine import by 50% (Fig. 1b). These findings reveal that administration of the creatine mimetic RGX-202 significantly suppresses creatine import into cardiac and tumor tissues *in vivo*.

**Fig. 1.**
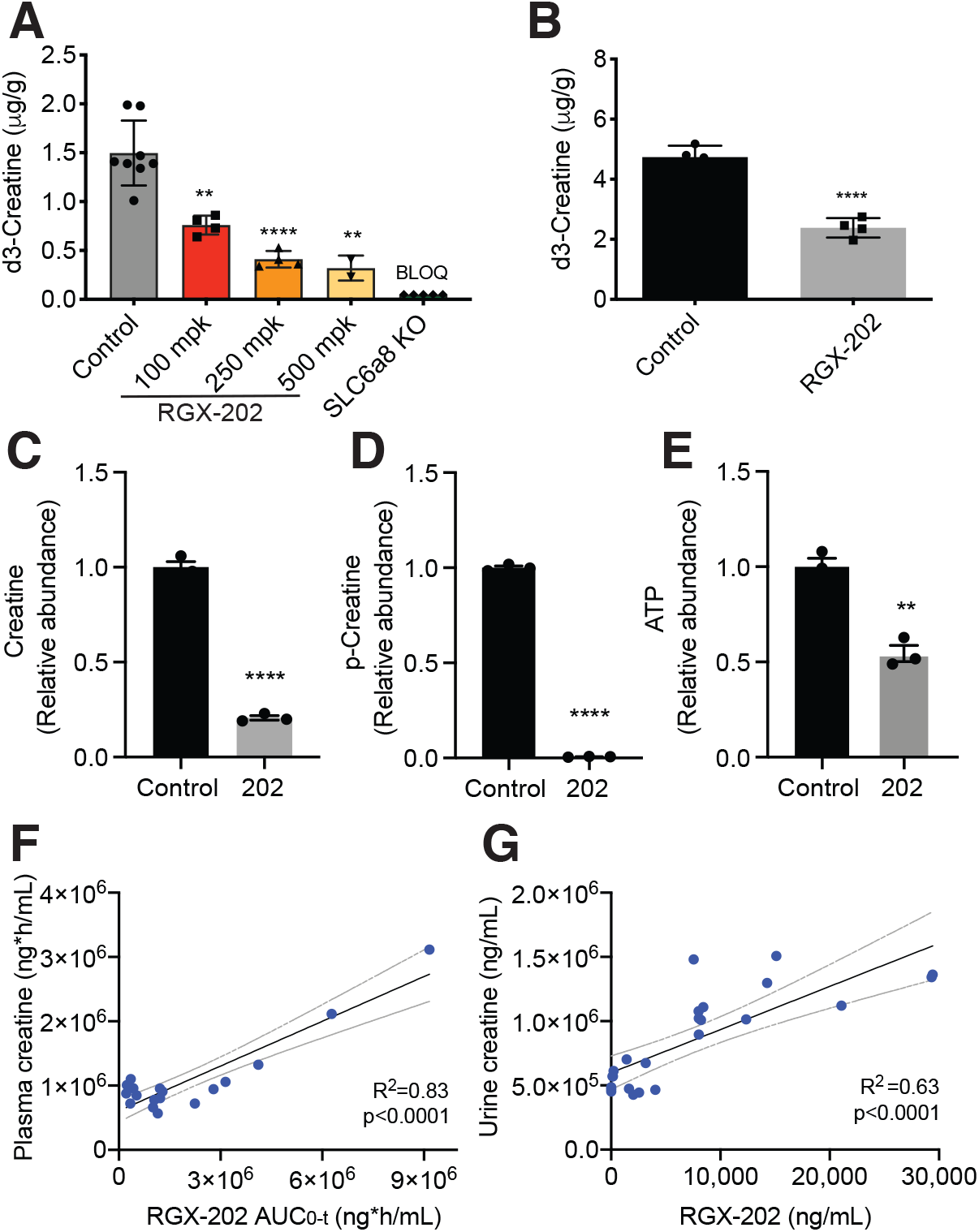
RGX-202 reduces cellular and tumoral creatine and phospho-creatine levels. **(A)** RGX-202 or vehicle control was injected (i.p.) into C57BL/6 wild type or SLC6A8 knock out mice, prior to injection of d3-creatine (i.p.). Mice were euthanized, hearts removed, and metabolites extracted for d3-creatine quantification by LC-MS/MS; n = 3-8 independent experiments; p = 0.0016 (100 mg/kg), p<0.0001 (250 mg/kg), p = 0.0014 (500 mg/kg). (**B)** UN-KPC-961 pancreatic tumor-bearing B6129SF1/J mice were fed a control or RGX-202 supplemented diet (800 mg/kg) for 35 days as soon as tumors reached ∼80 mm^3^. Mice were injected with 1 mg/kg d3-creatine (i.p.), tumors were extracted 1.5h later and d3-creatine levels were quantified by LC-MS/MS; n = 4 per group, p<0.0001. **(C-E)** 3×10^5^ Lvm3b cells were treated with vehicle control or 10 mM RGX-202 and incubated under hypoxia (0.5% O2) for 24h. Metabolites were extracted and creatine (p<0.0001) **(C),** phosphocreatine (p<0.0001) **(D)** and ATP (p = 0.0017) **(E)** levels analyzed by LC-HRMS; n = 3; representative of three independent experiments. P values are based on two-sided t tests +/- SEM **(A-E)**. **(F, G)** Correlation analysis of plasma creatine and RGX-202 exposure (AUC_0-t_) **(F)** and average urine creatine and RGX- 202 concentrations **(G)** measured over 24h in mice receiving either a control or RGX-202-01- supplemented diet at 100, 400 or 1200 mg/kg for 10 days; n = 6 per dose group.

We next assessed the impact of creatine transport inhibition on cellular levels of creatine, phosphocreatine and ATP. LC-MS/MS analysis of the highly metastatic CRC cell line LS174T Lvm3b (Lvm3b) treated with RGX-202 under hypoxia revealed a nearly complete depletion of phosphocreatine (>99%), greater than 79% reduction in cellular creatine and a substantial (46%) reduction in intracellular ATP levels relative to control cells (Fig. 1c-e). These findings are consistent with earlier reports describing reduced tissue creatine and phosphocreatine levels observed in rodents treated with β-GPA (*24, 26*). They are also consistent with our previous findings, demonstrating that shRNA-mediated depletion of the SLC6A8 transporter reduced phosphocreatine import and reduced ATP levels in CRC cells under hypoxia (*15*). These findings support a model whereby RGX-202- induced depletion of intracellular creatine and phosphocreatine leads to a consequential depletion of ATP, a critical energetic metabolite. The impairment of both phosphocreatine and ATP energetic metabolites poses a barrier for growth and survival under hypoxia and energetic stress.

We next assessed the effects of RGX-202-mediated inhibition of SLC6A8 on circulating levels of creatine in mice. Animals that received increasing doses of RGX-202-01 in the diet (100, 400 and 1200 mg/kg) exhibited an RGX-202 exposure-dependent increase in circulating plasma creatine levels, demonstrating accumulation of plasma creatine upon blockade of SLC6A8 in tissues (Fig. 1f). We also observed a substantial exposure-dependent increase in urinary creatine levels in the same animals (Fig. 1g), consistent with inhibition of creatine import into tissues causing increased creatine in circulation and subsequent excretion in urine, thus reducing creatine levels available to tumors.

### SLC6A8 inhibition exhibits broad anti-tumor activity against diverse primary and metastatic CRCs

Primary tumors exhibit hypoxia, especially upon progression to larger sizes and must gain metabolic adaptations to the hypoxic microenvironment to overcome this key barrier (*19*). To determine whether SLC6A8 inhibition impacts growth of established CRC tumors, we implanted highly aggressive metastatic Lvm3b (KRAS G12D) cells subcutaneously into athymic mice and began RGX-202 treatment after the tumors became palpable (>30 mm^3^). Oral administration of RGX-202 caused a ∼50% tumor growth inhibition (Fig. 2a). Interestingly, the effects of RGX-202 emerged after tumors reached larger sizes (> 500 mm^3^), consistent with the previously demonstrated role of creatine metabolism and SLC6A8 in hypoxic survival (*15*). RGX-202 treated mice exhibited a significant improvement in survival, experiencing a doubling of median survival time from 23 to 48 days. One of the nine immunodeficient mice experienced a complete tumor regression response and the tumor remained undetectable >38 days after treatment termination (Fig. 2b). We also observed anti-tumor efficacy in HCT116 (KRAS G13D) as well as HT29 (KRAS wild-type) human CRC tumors upon oral RGX-202 administration (Fig. 2C-E). Similar to Lvm3b, treatment of HT29 tumors induced regressions in 7 out of 10 mice upon tumors reaching a large size (>900 mm^3^) and treatment substantially prolonged overall survival (Fig. 2d and 2e). To determine if SLC6A8 inhibition could suppress progression of murine CRC in an immunocompetent model, we implanted KRAS G12D mutant CT26 murine CRC cells into syngeneic mice. RGX-202 significantly inhibited CT26 tumor growth (Fig. 2f). To determine if this approach is effective in treating larger tumors, we treated murine KRAS wild-type MC38 CRC tumors after volumes reached ∼150 mm^3^. RGX-202 administration substantially inhibited tumor growth in this immunocompetent model (Fig. 2g). Drug treatment significantly enhanced *in vivo* tumor apoptosis in a dose-dependent manner as quantified by cleaved Caspase-3 immunohistochemistry (Fig. 2h and 2i). Depletion of the SLC6A8 transporter in CRC cells was previously shown to substantially reduce liver metastatic colonization (*15*). We thus sought to determine whether inhibition of SLC6A8 could inhibit CRC metastasis. Highly metastatic Lvm3b cells, which harbored a luciferase reporter gene, were injected into the portal circulation of NOD SCID mice via intra-splenic injections and metastatic colonization to the liver was quantified by bioluminescent imaging. RGX-202 treatment reduced liver metastatic colonization by 8-fold (Fig. 2j and Supplementary Fig. 1a). To determine if the effect of SLC6A8 inhibition is cell-autonomous, we pre-treated G12D mutant PANC1 human pancreatic cancer cells *in vitro* with RGX-202 for 48h and subsequently injected these cells into the spleens of NSG mice. Pre-treatment with RGX-202 reduced pancreatic cancer liver metastatic colonization by 5-fold (Supplementary Fig. 1b and 1c), demonstrating that depleting cellular creatine/phosphocreatine levels via SLC6A8 inhibition significantly inhibits liver metastasis in a tumor cell-autonomous manner.

**Fig. 2.**
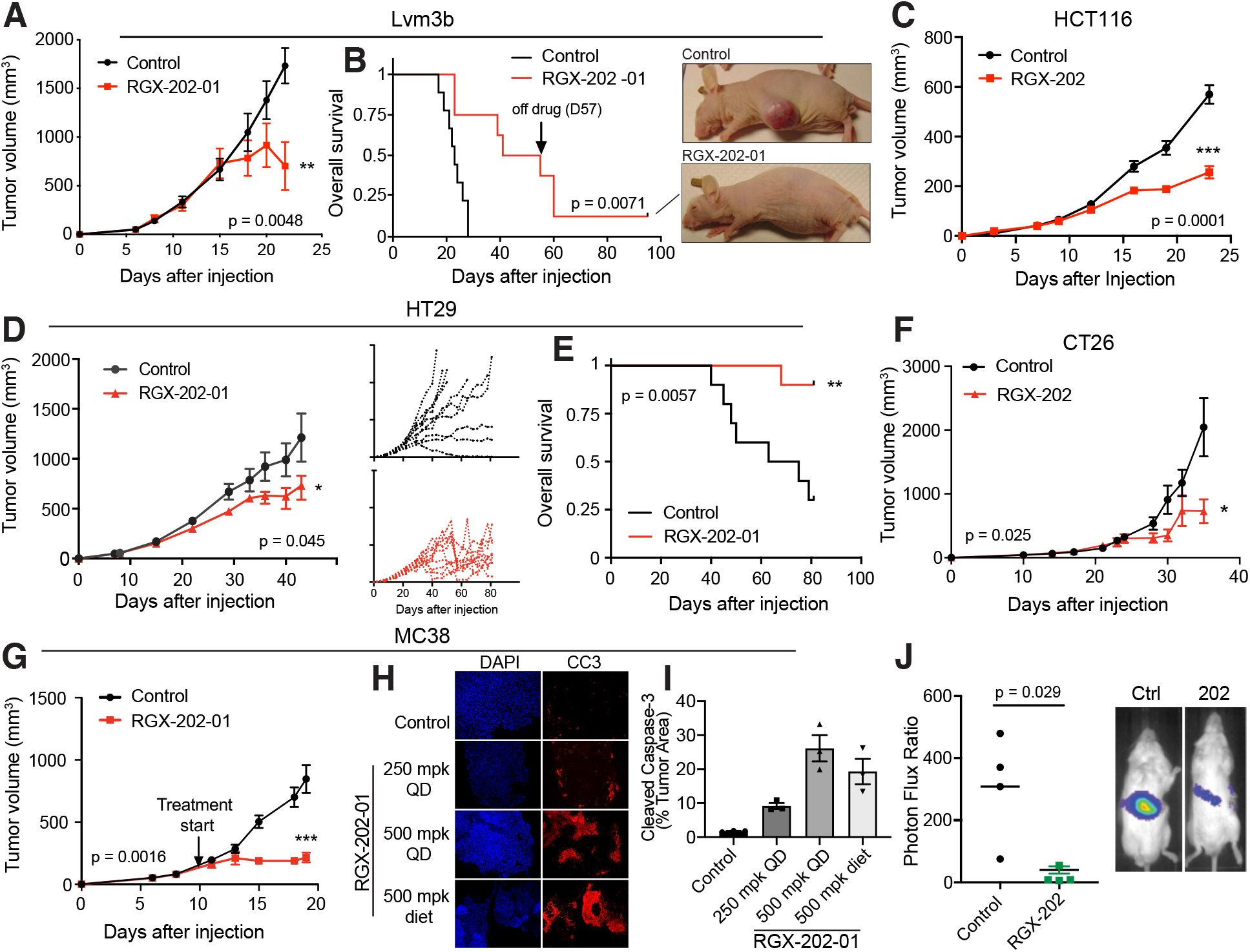
SLC6A8 inhibition exhibits anti-tumor activity against primary and metastatic CRCs of different backgrounds. **(A, B)** Subcutaneous tumor growth **(a)** and Kaplan-Meier survival curves **(B)** by 1 x 10^6^ Lvm3b cells in athymic nude mice. Daily oral gavage of RGX-202-01 (200 mg/kg) started at a tumor size of ∼50 mm^3^, n = 8-9 per group. Pictures show a control and RGX-202-01-treated mouse. **(C)** Subcutaneous tumor growth by 0.5 x 10^6^ HCT116 cells in athymic nude mice, receiving a control diet or RGX-202- supplemented diet (800 mg/kg) as soon as tumors were palpable; n = 5 per group. **(D, E)** Subcutaneous tumor growth **(D)** and Kaplan-Meier survival curves **(E)** by 2 x 10^6^ HT29 cells in athymic nude mice receiving a control diet or RGX-202-01-supplemented diet (800 mg/kg) when tumors reached ∼55 mm^3^; insets represent growth curves for individual tumors; n = 10 per group. **(F)** Subcutaneous tumor growth by 0.5 x 10^6^ CT26 cells in BALB/c mice receiving a control diet or RGX-202-01-supplemented diet (500 mg/kg) starting at a tumor size of ∼100 mm^3^; n = 7-8 per group. **(G)** Subcutaneous tumor growth by 0.5 x 10^6^ MC38 cells in C57BL/6 mice. Mice received a control diet or RGX-202-01-supplemented diet (500 mg/kg) starting at a tumor size of ∼150 mm^3^; n = 7 per group. **(H, I)** MC38 tumors treated with RGX-202-01 by oral gavage (QD 250 mg/kg p = 0.0011; 500 mg/kg p = 0.0032) or through diet (diet, p = 0.0092) for 9 days were extracted and immunostained for cleaved-Caspase 3 (CC3). Quantification **(I)** of the percentage of tumor area stained with cleaved-Caspase 3; n = 4 per group +/- SEM. **(J)** Bioluminescence plot of liver colonization by 5×10^5^ human Lvm3b metastatic colon cancer cells after intra-splenic injections into NOD-SCID mice. Mice were imaged on day 14 after injection; n = 4 per group. P values are based on two-sided t tests **(A,C,D, F-I)**, Log-rank Mantel-Cox test **(B,E)** or Mann-Whitney test **(J)**.

### SLC6A8 inhibition exhibits broad activity against CRC PDX models

Patient derived xenografts (PDXs) are thought to better recapitulate human tumor pathology and clinical drug responses (*27–30*). Therefore, we assessed the efficacy of SLC6A8 therapeutic inhibition on the growth of established (100-250 mm^3^) human KRAS wild-type and KRAS mutant CRC PDX tumors. Oral RGX-202 administration caused a 65% reduction in the growth of the CLR4 KRAS wild-type PDX model (Fig. 3a). Moreover, treatment also elicited regression responses in three PDX models harboring distinct KRAS mutations (Fig. 3b to 3d; CLR7 KRAS G12V, CLR24 KRAS G12C, and CLR30 KRAS G12R). We next sought to address patient-to-patient heterogeneity and predict potential clinical-trial responses by conducting a mouse PDX trial based on a 1×1 trial design where two mice become inoculated with the same PDX model and one mouse receives RGX-202 and the other control treatment (*31*). Such a trial design was developed to mimic a patient clinical trial and has shown reproducibility and translatability of therapeutic responses (*31*). We included 43 well documented colorectal cancer PDXs of various mutational backgrounds including both KRAS wildtype and KRAS mutant subtypes (Supplementary Table 1). Of these models, 49% of tumors harboring diverse KRAS mutant subtypes exhibited single agent anti-tumor efficacy greater than 30% relative to matched control animals (Fig. 3e). RGX-202 treatment also demonstrated anti-tumor efficacy in 30% of BRAF mutant tumors (2/7) (Supplementary Table 1). These findings reveal that SLC6A8 inhibition by oral RGX-202 treatment exhibits single agent anti-tumor efficacy against a broad range of CRC tumor subtypes and suggests potential for clinical benefit in both KRAS wild-type and KRAS mutant cancers.

**Fig. 3.**
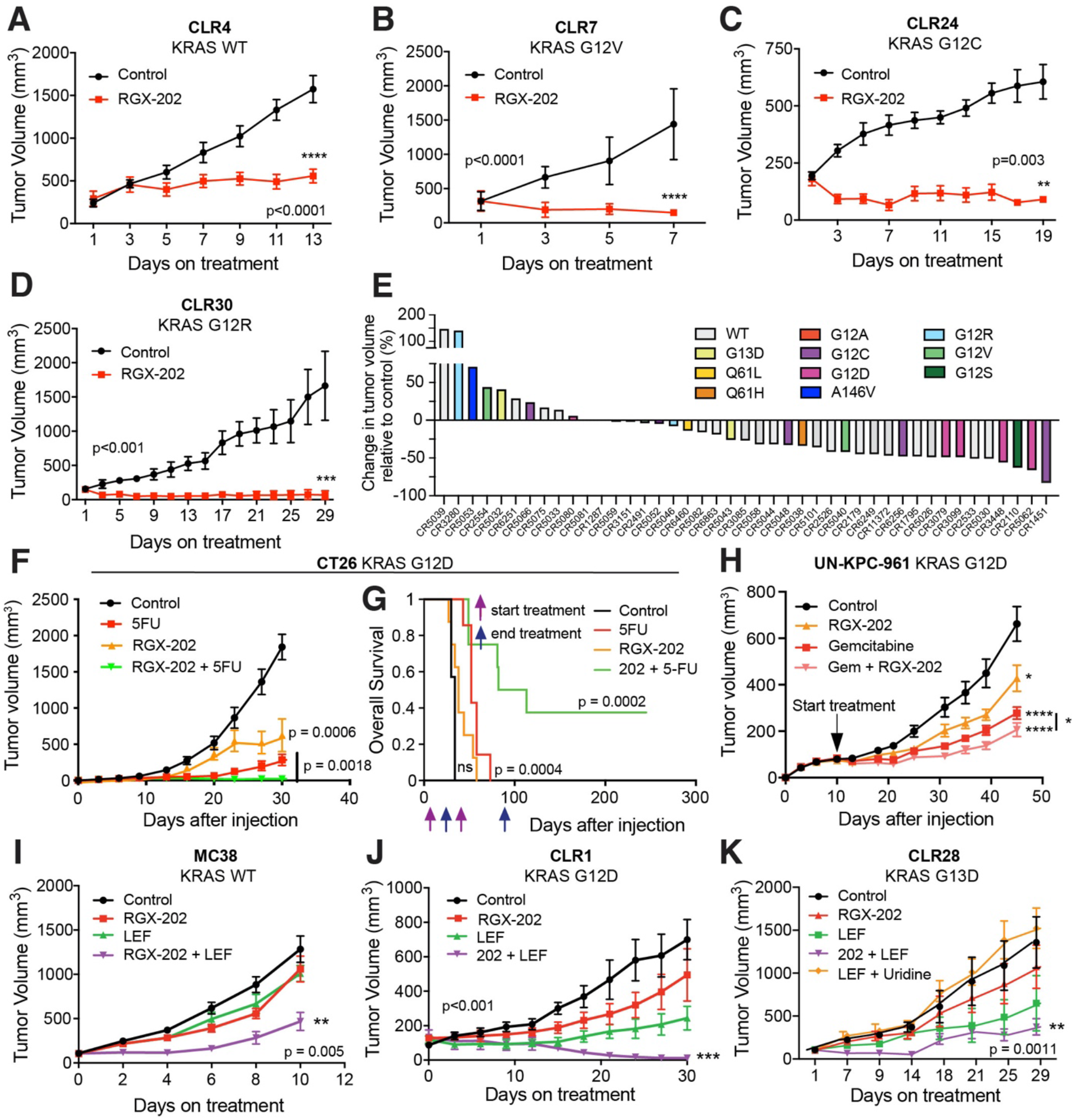
RGX-202 inhibits growth of KRAS wild-type and mutant CRC PDX and synergizes with 5-FU and leflunomide. **(A-D)** Subcutaneous tumor growth by ∼25 mm^3^ PDX fragments implanted in athymic nude mice receiving a control or RGX-202-supplemented diet (800 mg/kg) starting at a tumor size of ∼100 mm^3^ (CLR4), 250 mm^3^ (CLR7), 200 mm^3^ (CLR24) and 150 mm^3^ (CLR30); n = 10 per group. **e,** Waterfall plot of responses to RGX-202-01 across 43 colorectal PDX models; each bar represents an individual PDX and colors represent KRAS mutations. Mice were treated with control or RGX-202-01- supplemented diet at ∼400 mg/kg for 21 days. (**F, G)** Subcutaneous tumor growth **(F)** and Kaplan-Meier survival curves **(G)** of 1.5 x 10^5^ CT26 cells in BALB/c mice treated with control, RGX-202- formulated diet (800 mg/kg, p=0.0006), 5-FU (50 mg/kg/week) or a combination of RGX-202 and 5- FU; n = 7-8 per group. (**H)** Subcutaneous tumor growth of 2.5 x 10^6^ UN-KPC-961 cells in B6129SF1/J mice receiving control, RGX-202-formulated diet (800 mg/kg, p = 0.021), gemcitabine (i.p. 100 mg/kg, p<0.0001) or a combination of RGX-202 and gemcitabine (p<0.0001); n = 10 per group, *p (combination) = 0.04, one-tailed t test. **i,** Subcutaneous tumor growth of 1×10^6^ MC38 cells in C57BL/6 mice receiving a control diet, RGX-202-supplemented diet (200 mg/kg), leflunomide (2.5 mg/kg) or a combination of RGX-202 and leflunomide; n = 10 per group. **(J)** Subcutaneous tumor growth by CLR1 PDX fragments implanted into athymic nude mice. Treatment with a control diet, a diet supplemented with 800 mg/kg RGX-202, leflunomide (7.5 mg/kg) or a combination of RGX-202 and leflunomide started when tumors reached ∼100 mm^3^, n = 6 per group. (**K)** Subcutaneous tumor growth by CLR28 PDX fragments in athymic nude mice. Treatment with a control diet, RGX-202-supplemented diet (800 mg/kg), leflunomide (7.5 mg/kg), a combination of RGX-202 and leflunomide or uridine (1g/kg) and leflunomide started at ∼100 mm^3^; n = 6 per group. P values are based on two-sided t tests **(A-D,F,H-K)** or Log-rank Mantel-Cox test **(G)**.

### SLC6A8 inhibition synergizes with 5-FU and oral leflunomide therapy

We next determined whether SLC6A8 inhibition could cooperate or synergize with other therapeutics. 5-FU is the backbone of many chemotherapeutic regimens used in CRC. We established palpable CT26 subcutaneous tumors in immunocompetent syngeneic mice and began treatment with vehicle control, oral RGX-202, 5-FU, or a combination regimen comprising RGX-202 and 5-FU. While single-agent RGX-202 or 5-FU significantly suppressed tumor growth (66% inhibition RGX-202, 85% inhibition 5-FU), combined RGX-202/5-FU caused a 99% tumor growth reduction (Fig. 3f).

Importantly, survival studies revealed that combined RGX-202/5-FU therapy dramatically enhanced survival of mice, causing 40% of mice to experience complete regression responses and long-term (>240 days) survival (Fig. 3g) despite cessation of treatment at day 85, 160 days prior to termination of the study. RGX-202 also elicited enhanced anti-tumor efficacy in combination with 5-FU and irinotecan, a standard of care regimen for metastatic colorectal cancer (Supplementary Fig. 2). We also observed that combined treatment of established UN-KPC-961 (Kras^G12D^;Trp53^R172H^) pancreatic cancer tumors with RGX-202 and gemcitabine, a deoxycytidine analog and approved pancreatic cancer therapeutic, elicited greater tumor suppression relative to either gemcitabine or RGX-202 alone, though the enhanced effects were modest compared to combination effects observed in CRC (Fig. 3h).

We had previously shown that under hypoxia, metastatic CRC tumors induce pyrimidine nucleotide biosynthetic pathways and become sensitized to inhibition of the dihydroorotate dehydrogenase (DHODH) enzyme, which catalyzes a critical step during pyrimidine nucleotide biosynthesis (*22, 32*). Treatment of mice with the oral DHODH inhibitor leflunomide, which is widely used as a rheumatoid arthritis drug, inhibited CRC primary tumor and metastatic progression (*22*). Combined treatment of established MC38 KRAS wild-type syngeneic CRC tumors with RGX-202/leflunomide elicited synergistic activity relative to single-agent treatment with either drug (Fig. 3i). We observed similar examples of synergy in PDX models. For example, the CLR1 KRAS G12D mutant and CLR28 KRAS G13D mutant PDXs exhibited tumor regressions upon combined administration of RGX- 202/leflunomide relative to tumor growth suppression responses by either agent alone (Fig. 3j and 3k). To determine if the effect of leflunomide on tumor growth repression in the CLR28 model was secondary to pyrimidine nucleotide depletion, we included a cohort that received leflunomide and Uridine, a pyrimidine nucleotide. Uridine supplementation rescued tumor growth inhibition by leflunomide, consistent with pyrimidine nucleotide levels being limiting for tumor growth (Fig. 3k).

Overall, these findings reveal that RGX-202 can cooperate or synergize with the standard of care agent 5-FU as well as the FDA approved rheumatologic oral drug leflunomide.

### Tumoral CKB expression as a predictive biomarker of SLC6A8 inhibition response

We next sought to identify a predictive biomarker for therapeutic response to SLC6A8 inhibition by RGX-202. Our observations of therapeutic efficacy upon SLC6A8 inhibition in both KRAS mutant, KRAS wildtype, and mismatch repair mutant CRC lines suggested that sensitivity to SLC6A8 inhibition was not solely attributable to oncogenic or genomic instability mutational backgrounds (Supplementary Table 2). We previously showed that the CKB enzyme is released from CRC cells and acts upstream of the SLC6A8 transporter by generating the energetic metabolite phosphocreatine, which is imported through the SLC6A8 transporter (*15*). Extracellular phosphocreatine supplementation rescued hypoxic survival impairment caused by CKB depletion in an SLC6A8- dependent manner (*15*). CKB expression was also shown to be repressed by microRNA-mediated silencing, raising the possibility that variation in its expression may predict sensitivity to therapeutic targeting of this axis (*15*). We therefore tested a collection of gastrointestinal cell lines as xenografts for responsiveness to RGX-202 (Supplementary Table 2 for details on the cell lines). An independent cohort of animals were simultaneously inoculated with the same xenografts and tumors were allowed to reach ∼ 500 mm^3^, at which point tumoral SLC6A8 and CKB gene expression by qPCR and CKB protein expression by immunohistochemistry (IHC) were assessed (Fig. 4a). SLC6A8 was not tested by IHC due to lack of an SLC6A8-specific antibody. Included in this assessment were tumors that responded to treatment and those that did not (Fig. 4b-d, Supplementary Table 2 and Supplementary Fig. 3). We observed that therapeutic responses to RGX-202 positively correlated with increased CKB mRNA expression (Fig. 4e) but not SLC6A8 mRNA expression (Supplementary Fig. 4a). Consistent with this, the extent of CKB protein expression—as assessed by Tumor Proportion Score (TPS) —also positively correlated with the magnitude of therapeutic response (Fig. 4f). We observed that tumors expressing elevated CKB protein levels (Fig. 4b) exhibited greater tumor growth repression responses relative to those expressing reduced CKB levels (Fig. 4e, 4f and Supplementary Fig. 4b-c). To determine if this observation could be recapitulated in an independent dataset, we assessed the association of CKB protein expression with therapeutic responses in the aforementioned mouse RGX- 202 1×1 PDX trial. We observed a similar predictive association of elevated CKB protein expression by IHC with anti-tumor response (Fig. 4g). These results demonstrate that elevated tumoral CKB mRNA and protein expression predict for enhanced responsiveness to SLC6A8 inhibition by RGX- 202, consistent with CKB being an upstream component of the CKB/SLC6A8 phosphocreatine metabolic axis in CRC and a gene that exhibits post-transcriptional expression modulation and variation in cancer. To assess what fraction of patients with metastatic CRC harbor tumors with elevated CKB protein expression, we immunohistochemically stained 23 human metastatic CRC specimens for CKB. We observed CKB positivity (TPS >5%) in the majority (56%;13/23) of the tumor tissue samples (Supplementary Fig. 4 d, e). Overall, these findings reveal CKB as a potential patient stratification biomarker for clinical development of SLC6A8 inhibitors.

**Fig. 4.**
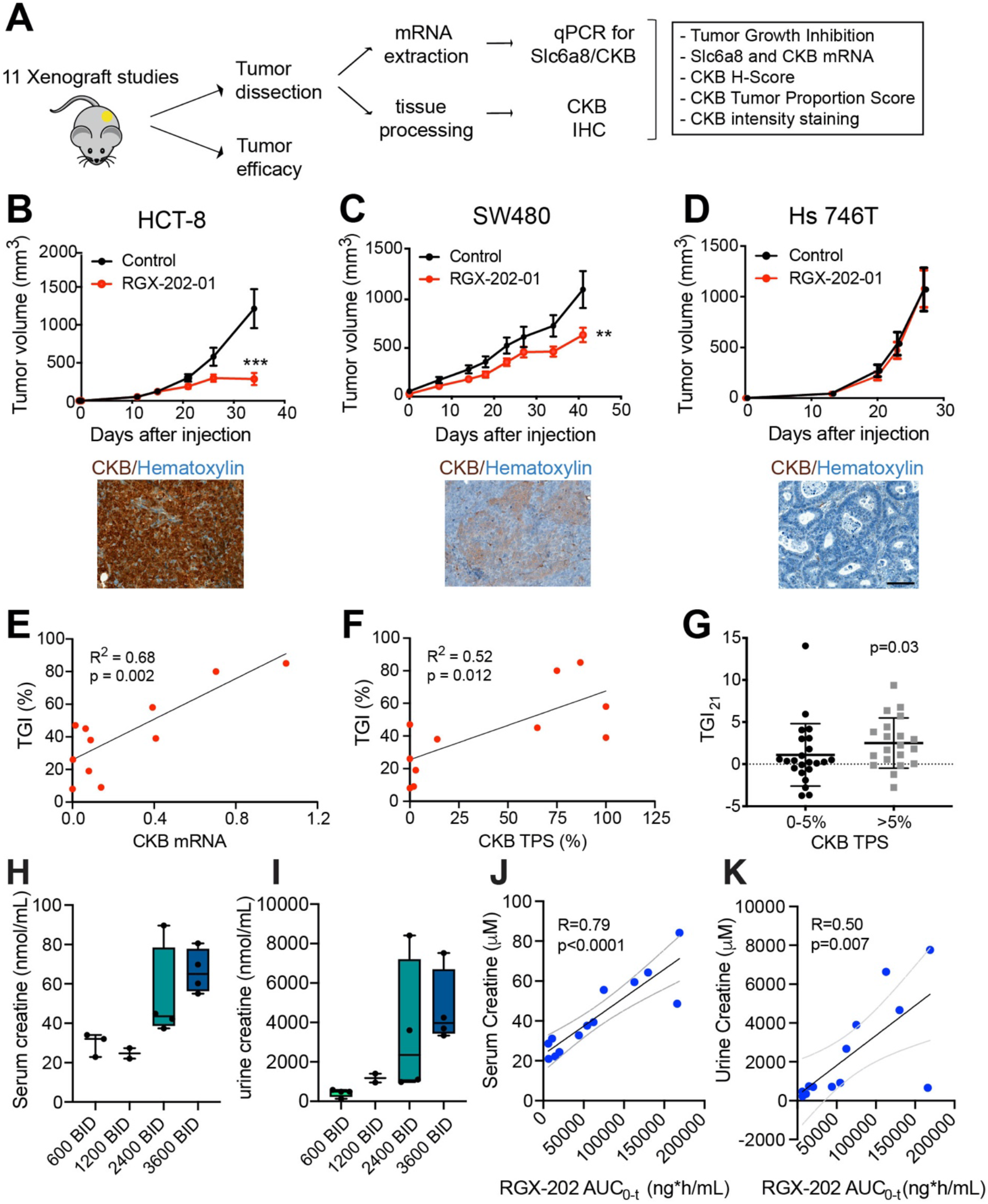
Tumoral CKB expression and creatine levels as predictive and pharmacodynamic biomarkers of SLC6A8 inhibition. **(A-D)** Schematic showing the experimental design **(A)**. Tumor growth by 5×10^6^ HCT-8 (p = 0.0004) **(B)**, 5×10^6^ SW480 (p = 0.01) **(C)** and 2×10^6^ Hs746T **(D)** cells subcutaneously injected into athymic nude mice receiving a control diet or RGX-202-01-supplemented diet (∼800 mg/kg). Pictures show IHC of CKB expression (brown), counterstained with Hematoxylin (blue) in the control tumors; n = 8- 10 per group. P based on two-sided t tests. **(E, F)** Linear regression analyses of tumor growth inhibition (TGI) and CKB mRNA **(E)** and CKB tumor proportion score (TPS) **(F)**; n = 11 models. (**G**) Non-parametric analysis of tumor growth inhibition on Day 21 relative to control from the 43 PDX models, stratified into models with low (0-5%) or >5% CKB TPS. **(H, I)** Box-and-whisker plot representing the median of creatine concentrations in the serum (**H**) and urine (**I**) measured at 4h (serum) and 12h (urine) after RGX-202 administration; n = 2-4 patients per group. (**J,K)** Correlation analysis of serum (**J**) or urine (**K**) creatine concentration and RGX-202 blood exposure (AUC0-t). Dashed lines denote 95% confidence interval. Patients were sampled on Cycle 1 Day 15, n = 13 patients total **(H-K)**.

### RGX-202 increases serum and urinary creatine excretion in patients with cancer

Motivated by the robust pre-clinical efficacy of RGX-202, we initiated a multicenter Phase Ia/b clinical trial in patients with advanced gastrointestinal cancers that had progressed on standard of care regimens (ClinicalTrials.gov, NCT03597581). In the Phase 1a dose escalation stage of the study, patients received oral RGX-202 treatment at doses ranging from 600mg to 3600mg twice daily, on a continuous regimen. Blood and urine samples were collected from 13 patients on Day 15 during the first cycle (28 days), and bioanalytic analyses were conducted by an independent contract research laboratory using commercially available proprietary assays. Consistent with our observations in mice where RGX-202-mediated suppression of creatine uptake in tumors and tissues led to increased circulating and urinary excretion levels of creatine, we observed that patient serum and urine creatine levels increased upon RGX-202 administration in cancer patients (Fig. 4h and i). Creatine concentrations in both serum and urine showed positive correlations with systemic exposure to RGX- 202 (Fig. 4j and 4k). These findings provide proof-of-concept for therapeutic targeting of creatine metabolism in humans, mirroring our experimental observations in mice.

## Discussion

The creatine/phosphocreatine bioenergetic shuttle is a critical system that allows highly metabolic tissues to respond to energetic stress quickly through the generation of high energy ATP in a reaction that does not require oxygen. CRC and pancreatic cancers are particularly hypoxic malignancies (*19–21*). The ability of such cancers to over-express and release CKB as a means of generating extracellular phosphocreatine for import through the SLC6A8 transporter enables cancer cells to enhance availability of high energy phosphate for ATP generation in the context of hypoxic and metabolic stresses. SLC6A8 inhibition by RGX-202 reduced tumor growth in numerous syngeneic, xenograft and PDX mouse models and also suppressed liver metastasis. Anti-tumor efficacy was associated with enhanced tumor apoptosis, consistent with prior findings supporting a critical role for creatine metabolism in CRC progression and hypoxic survival (*15*). Our work reveals that mutational background is not a significant predictor of response to SLC6A8 inhibition and that RGX-202 impairs the growth of a broad set of CRC tumors of distinct KRAS and mismatch repair mutant backgrounds suggesting that rather diverse CRC tumors exploit creatine/phosphocreatine metabolism to drive progression. Consistent with a critical role for CKB in this pathway, we find that CKB expression levels are associated with response to RGX-202. Tumors with high CKB expression responded better than those with reduced levels, suggesting that CKB over-expressing tumors are more dependent on phosphocreatine as an energy source and thus more sensitive to its inhibition. These findings support the potential for assessing patient tumor CKB expression as a molecular biomarker for patient stratification in clinical trials.

Our data also reveal that inhibition of SLC6A8 and blockade of cellular creatine/phosphocreatine uptake leads to increased creatine excretion in the urine, both in mice and in patients. Urinary creatine levels positively associated with blood RGX-202 concentrations and represent a direct measure of SLC6A8 inhibition. The opportunity for non-invasive monitoring of the pharmacodynamics of creatine transport inhibition through urinalysis of creatine and potentially additional metabolites, provides a simple and rapid method for assessing target engagement in patients.

Our findings reveal both single-agent activity and combination efficacy of RGX-202 in CRC. Because 5-FU is the mainstay chemotherapeutic used in CRC, we tested the activity of RGX-202 in combination with 5-FU and 5-FU/irinotecan. Our data show synergistic effects of SLC6A8 inhibition and complete tumor regressions with these standard of care regiments. Similarly, we observed robust anti-tumor efficacy when combining RGX-202 with the DHODH inhibitor leflunomide, an oral therapy that was previously shown to suppress CRC growth under hypoxia (*22*). The synergistic effects observed likely reflect the impairment of these two orthogonal metabolic pathways— creatine/phosphocreatine and pyrimidine biosynthesis—that appear critical for the growth and survival of colorectal cancer cells under hypoxia. Collectively, these results provide a rationale for incorporating RGX-202 into combination regimens including standard of care agents or those targeting nucleotide synthesis. Future studies could investigate the impact of this therapy in the adjuvant setting, as a means of suppressing CRC metastatic progression.

Overall, this work supports the potential for therapeutic targeting of creatine metabolism in CRC through the inhibition of the SLC6A8 transporter. While we have demonstrated pharmacodynamic effects on creatine metabolism in patients upon RGX-202 treatment, ongoing trials are testing the efficacy of this therapeutic approach in patients. To this end, this treatment approach is currently being tested in a multi-center national Phase 1b/2 clinical trial in CRC (ClinicalTrials.gov, NCT03597581).

## Materials and Methods

### Experimental design

The number of samples for each group was chosen based on knowledge on intra-group variation and expected effect size. Sample sizes for *in vitro* experiments were chosen based on prior knowledge on intra-group variation. Data was collected until reaching a pre-determined end-point (in vitro assays, PDX trial and clinical sample analysis) or tumor burden exceeded >1500 mm^3^. No data were excluded. Replications were performed as noted in text and figure legends. Replications for all presented experiments were successful. Samples were allocated randomly if possible. Animals were randomized prior to start treatment and mice were sex- and age-matched. No blinding was performed in the in vivo experiments due to cage labeling requirements. Quantification of CKB signal by IHC was blinded.

### Animal strains

All mouse experiments and procedures were approved by the Institutional Animal Care and Use Committees (IACUC) at the New York Blood Center (NYBC), The Rockefeller University, Memorial Sloan Kettering Cancer Center and Crown Biosciences. C57BL/6 (JAX stock #000664, RRID: IMSR_JAX:000664), NOD-SCID (JAX stock #001303), Athymic nude (J:NU, JAX stock #007850), NOD *scid* gamma (NOD.Cg-Prkdcscid Il2rgtm1Wjl/SzJ, JAX stock #005557) and B6129SF1/J (JAX stock #101043) mice were purchased from the Jackson Laboratory. BALB/c mice (stock #028) were purchased from Charles River. SLC6A8 knockout mice were originally obtained from the laboratory of Dr. Skelton (*33*) and bred in-house. BALB/c nude mice were purchased from Beijing Anikeeper Biotech Co.,Ltd (Beijing, China).

### Primary tumor growth studies

For primary tumor growth experiments, cells (suspended in 50 μl of PBS) were mixed 1:1 with Matrigel (356231, BD Biosciences, Bedford, MA) and subcutaneously injected unilaterally or bilaterally into the lower flank of 6 to 8-week old sex matched mice. Upon detection of tumor volumes reaching the size indicated in each figure, mice were randomly assigned to a drug treatment or a control cohort. RGX-202 was administered through formulated drug chow (Purina 5001, Research Diet, New Brunswick, NJ) at indicated doses or formulated in sterile drinking water for oral gavage.

Control cohorts received either regular chow (Purina 5001) or vehicle control. Where xenograft models were tested for biomarker analysis, 10-15 animals were inoculated with the indicated cell lines. Out of these animals, 7-10 mice were assigned to the efficacy portion and were treated for the duration indicated in each figure. The remaining 3-5 animals were sacrificed when tumors reached ∼500 mm^3^.

Protein and RNA analysis of CKB expression was conducted on the control tumors. Tumor growth measurements were taken using a digital caliper on the days indicated throughout the course of the experiment. Tumor volume was calculated using the formula: volume = (longest diameter)/2 x (shortest diameter)^2^. For survival analysis, mice were euthanized when total tumor burden approached IACUC guidelines with a tumor burden exceeding 2000 mm^3^ in volume. Tumor growth inhibition (TGI) was calculated using the formula TGI (%) = (((ΔC_i_-ΔC_0_) – (ΔT_i_-ΔT_0_))/(ΔC_i_-ΔC_0_))*100%.

### Patient-derived xenograft (PDX) studies

Implantation of PDX’s was conducted as described before (*22*). Briefly, 20-30 mm^3^ tumor fragments were subcutaneously implanted bilaterally into the flank of 6 to 8-week old age-matched athymic nude mice under general anesthesia with 100 mg/kg ketamine (995-2949-100 mg/mL, Henry Schein Animal Health, Melville, NY) and 10 mg/kg xylazine (X1251, Sigma Aldrich, St. Louis, MO).

### Metastasis assays

Experiments were conducted by intra-splenic injection of 5 x 10^5^ luciferase-labeled Lvm3b cells suspended in 50 μL PBS into 6 to 8-week-old NOD SCID mice that were anesthetized through i.p. injection of ketamine/xylazine solution (100 mg/kg Ketamine, 10 mg/kg xylazine). The day after tumor cell inoculation, mice were randomly assigned to a control or RGX-202 treatment group.

Control mice received i.p. injections of 200 µL of PBS and treatment mice received 200 µL of 0.5 M RGX-202 (∼650 mg/kg). Treatment continued daily until the end of the experiment. In the PANC1 experiment, cells grown in D10F complete medium were pre-treated in vitro with or without RGX-202 at a dose of 10 mM (1.31 mg/mL) RGX-202 for 48 hours prior to injection into the mice. There was no treatment of the mice after the cells were injected. Bioluminescence measurements were conducted once per week for the duration of the experiment. One hundred µL of *D*-Luciferin (88292, Thermofisher, Waltham, MA, 1 g in 60 mL sterile DPBS,) was injected into the venous sinus and the bioluminescence signal over the liver was measured using an IVIS Spectrum In Vivo Imaging System (Perkin Elmer). Photon flux ratio is the ratio of bioluminescence signal at a given time point to the signal on day 0.

### Drug treatments

5-Fluorouracil (5-FU) (F6627, Millipore Sigma, St. Louis, MO) was administered in sterile 0.9% NaCl by intraperitoneal injections (i.p.) once per week as indicated in the figures. Gemcitabine (G6423, Millipore Sigma, St. Louise, MO) was administered at 100 mg/kg/week i.p. in PBS. Irinotecan (I1406- 50mg, Millipore Sigma, St. Louise, MO) was administered at 15 mg/kg/week i.p. in 2.5% DMSO 97.5% NaCl 0.9%. Leflunomide was formulated in DMSO and administered by daily i.p. injections at 2.5 or 7.5 mg/kg at 0.5 mL/kg. Uridine was formulated in sterile water at 800 mg/kg and administered by daily i.p. injections.

### Cell culture

UN-KPC-961 cells were obtained from Professor S.K. Batra at Eppley Institute for Research in Cancer (Omaha, Nebraska). COLO 205 (CCL-222), HCT-15 (CCL-225), HT-29 (HTB-38), SW 480 (CCL- 228), HCT 116 (CCL-247), HCT-8 (CCL-244), NCI-H508 (CCL-253), HepG2 (HB-8065), Hs746T (HTB-135), CT26 (CRL-2638), LS-174T (CL-88), PANC1 (CRL-1469) and NCI-N87 (CRL-5822) cell lines were purchased from ATCC (Baltimore, MD) and maintained in standard conditions following manufacturer’s instructions. Lvm3b was generated by in vivo selection from the parental cell line LS-174T (ATCC, Baltimore, MD) (*15*). UN-KPC-961 cells were maintained in DMEM (11960- 044, Gibco, Langley, OK), 7.5% Sodium Bicarbonate (25080-094, Gibco, Langley, OK), 1% Penicillin-Streptomycin (Lonza, 17-745E), 10% Fetal Bovine Sera (F4135, Sigma, St. Louis, MO), 200 mM L-Glutamine (25030081, Gibco, Langley, OK), 1 mM HEPES (15630080, Gibco, Langley, OK) and 50 mg/ml Gentamicin (15750078, Gibco, Langley, OK). MC38 cells were cultured in DMEM, 10% Fetal Bovine Serum and 1 mM HEPES.

### 1×1 PDX trial (HuTrial)

Details on the models are found in Supplementary Table 1. PDX models were inoculated with a PDX cell suspension or a tumor fragment as follows. Cryogenic vials containing PDX tumor cells were thawed and cells washed in RPMI, counted, and resuspended in cold RPMI at a concentration of 50,000-100,000 viable cells/50 mL. Cell suspensions were mixed with an equal volume of Cultrex ECM. 100 1L of the cell suspension in ECM media was subcutaneously injected into the rear flank of 5 female NOD-SCID mice per model. Alternatively, tumor fragments from stock mice were harvested and used for inoculation into mice. Five 6 to 8-week old female Balb/C nude mice per model was inoculated subcutaneously in the right flank with a primary human tumor fragment (2-3 mm in diameter) for tumor development. Two out of these five mice were randomized into 2 groups (1 mouse per group) when average tumor volume reached 100-150 mm^3^ and treatment with either control chow (Purina 5001) or RGX-202-formulated chow (in Purina 5001, Research Diet, New Brunswick, NJ) at ∼400 mg/kg started within 24h after randomization. Randomization was performed based on “Matched distribution” method (StudyDirector^TM^ software, version 3.1.399.19). Tumor growth inhibition on Day 21 (TGI_21_) was calculated using the formula TGI_21_ = ((C_21_-C_0_)/C_21_) – ((T_21_-T_0_)/T_0_). The PDX trial was conducted at Crown Biosciences (San Diego, CA and Taicang Jiangsu Province, China).

### Urine and plasma creatine analysis

Six to eight-week old female CD-1 mice were fed with control or RGX-202-01-formulated chow (Purina 5001, Research Diet, New Brunswick, NJ) at 100, 400 and 1200 mg/kg for 10 days. Bleeds were collected in EDTA-coated blood collection tubes (02-669-33, Fisher Scientific) and plasma separated upon centrifugation. Plasma and urine samples were collected from both the control and RGX-202-treated cohorts on Day 10 (240 h, 246.5 h, 254.5 h) and Day 11 (264 h) timepoints. Samples were flash frozen and stored at −80°C until bioanalysis for RGX-202 and creatine levels at Pharmaron (Beijing, China). Mice plasma and urine analysis were conducted as a non-GLP study according to standard operating procedures at Pharmaron (Beijing, China). Briefly, concentration of RGX-202-001 was determined using a surrogate analyte RGX-202-^13^C1^15^N_2_ by liquid chromatography tandem mass spectrometry (LC-MS/MS). Lower limit of quantification (LLOQ) was determined to be 50.0 ng/mL. The in-life portion of this study was conducted at AJES LifeSciences, LLC/L2P.

### Creatine-(methyl-d3) uptake in the tumors and heart

Three to four B6129SF1/J male mice harboring UN-KPC-961 tumors that were treated with either control chow or RGX-202-formulated chow at ∼800 mg/kg for 35 days, received one dose of 1 mg/kg creatine-(methyl-d3) (DLM-1302-0.25, Cambridge Isotope Laboratories, Tewksbury, MA) by i.p. injection. One and a half hours later, the animals were euthanized, the tumors and hearts were extracted, flash frozen in liquid nitrogen and submitted to Seventh Wave Laboratories for bioanalysis. Three to four 6-9-week-old C57BL/6J male wild-type male mice received one dose of RGX-202 at 100, 250 or 500 mg/kg, respectively, in sterile 0.9% NaCl. Seven minutes following injection of RGX- 202 or vehicle control, one dose of d3-creatine was administered at 1 mg/kg in sterile 0.9% NaCl by i.p. injection. SLC6A8 knock-out mice received one dose of d3-creatine. One hour later, the mice were anesthetized using 2.5% isoflurane and the animal was perfused with 5-10 mL of DPBS that was injected into the left heart chamber. The heart was then extracted, and flash frozen in liquid nitrogen. Heart samples were stored at −80°C until being submitted to Seventh Wave Laboratories for analysis. Hearts and tumors were homogenized with a 70/30 methanol/water solution at a 3:1 (v:w) ratio. 20 μL of the homogenate was diluted with 1 μL of internal standard solution (Creatine-d5 in methanol). The samples were centrifuged, and the supernatant was analyzed. A fit-for-purpose, semi-quantitative liquid chromatography with tandem mass spectrometry (LC-MS/MS) method was developed at Seventh Wave Laboratories for the determination of d3-creatine and creatinine in heart and tumor tissue homogenate. Creatinine concentrations were used to normalize the signal of d3-creatine across samples.

### Real-time PCR analysis of CKB and SLC6A8 from tumor samples

RNA was extracted from 10 mg of flash frozen vehicle or RGX-202-treated tumors using Total RNA Purification kit (37500, Norgen Biotek, Thorold, Canada) according to the manufacturer’s instructions. RNase-Free DNase I (25710, Norgen Biotek, Thorold, Canada) was used to remove genomic DNA. cDNA synthesis was performed with 600ng of total RNA, using a Verso cDNA synthesis kit (AB- 1453/B, Fisher Scientific, Waltham, MA) according to protocol. qPCR was performed using TaqMan Fast Universal PCR Master Mix (2X), no AmpErase UNG (Fisher Scientific, Cat #: 4366073) and TaqMan probes (20X), with 1 µL of 1:5 diluted cDNA.

Predesigned Taqman Gene Expression Assays for CKB, SLC6A8 and GUSB (4331182, Fisher Scientific, Waltham, MA) were used to perform the reactions in a Step One plus Real-time PCR system (Applied Biosystems). GUSB was used as an internal control.

A standard curve for normalization was generated by preparing serial dilutions as follows. Undiluted cDNA from the HCT116 cell line at a stock concentration of 30 ng/µL was used to prepare dilutions of 12.5 ng, 4.17 ng, 1.25 ng, 0.42 ng, 0.125 ng and 0.013 ng. 1:5 diluted cDNA sample of HCT116 cell line was used as a reference that was run in all qPCR plates to account for plate to plate variability. All standards and samples were tested in quadruplicates.

### Histology, Immunohistochemistry and Immunofluorescence

Tumors were fixed overnight with 4%PFA (50-980-497, EMS, Hatfield, PA) at 4°C. After two washes with Phosphate Buffer Saline (PBS) (10010023, Gibco), tissue was embedded in paraffin following standard protocols. Tumors were sectioned using a microtome (Leica) and 5μm-thick sections were mounted on Superfrost Plus Microscope Slides (22-037-246, Fisher Scientific). The immunohistochemistry detection of CKB was performed at the Molecular Cytology Core Facility of Memorial Sloan Kettering Cancer Center, using Discovery XT processor (Ventana Medical System, Roche - Indianapolis, IN). The tissue sections were blocked first for 30 min in Background Blocking reagent (Innovex, catalog#: NB306). A rabbit monoclonal antibody (ab92452, Abcam) was used in 1 : 400 dilution. The incubation with the primary antibody was done for 5 hours, followed by 60 minutes with biotinylated goat anti-rabbit IgG (PK6101, Vector labs) in 5.75µg/ml, Blocker D, Streptavidin-HRP and DAB detection kit (Ventana Medical Systems) were used according to the manufacturer instructions. Slides were counterstained with hematoxylin and coverslipped with Permount (Fisher Scientific). Antibody specificity was determined by stainings conducted on HCT116 cells transiently transfected with CKB siRNAs. Immunofluorescence detection of cleaved Caspase-3 was performed as described in the above protocol with the following modifications: primary anti-cleaved Caspase-3 antibody (Asp175) (9664, Cell Signalling) was used at a 1:150 dilution. Incubation was performed overnight and after 3 washes with PBS, secondary antibody donkey-anti rabbit Alexa Fluor (A32795, Invitrogen) at a 1:600 dilution was incubated for 1h. Slides were mounted and counterstained with FluorSave reagent (345789, Sigma).

### Analysis of tumor samples

Six to eight tumors per xenograft model was assigned a score of 0-3 based on intensity of CKB staining. Score 0 is interpreted as negative for protein expression while scores 1, 2, and 3 are interpreted as positive staining for each core with 3 being maximal intensity. For each positive sample, the area (% of tumor) corresponding to each intensity staining was recorded to allow for calculation of percentage tumor positivity (Tumor Proportion Score). A weighted overall staining score (H-score) was calculated as (percentage area of 1+ staining x 1) + (percentage area of 2+ staining x 2) + (percentage area of 3+ staining x 3). Tumor sections were excluded from the analysis if that section was predominantly necrotic. In these cases, new tumor sections were stained to reach a minimum of n = 3-4 sections per tumor, 3 tumors per cohort.

### Metabolite extraction and liquid chromatography

Lvm3b cells were plated at 3 × 10^5^cells/well in triplicates in RPMI1640 in the presence of dialyzed FBS, 2mM glutamine and 6 mM glucose and allowed to adhere to the plate for 24 hr. Cells are treated with either control or 10mM RGX-202 for 24 hr in 0.5% O_2_. Cells were washed with ice cold 0.9% NaCl and harvested in ice cold 80:20 LC-MS methanol:water (*v/v*). Samples were vortexed vigorously and centrifuged at 20,000 *g* at maximum speed at 4°C for 10 min. The supernatant was transferred to new tubes. Samples were then dried to completion using a nitrogen dryer. All samples were reconstituted in 30 μl 2:1:1 LC-MS water:methanol:acetonitrile. The injection volume for polar metabolite analysis was 5 1l. Metabolite extraction and subsequent Liquid-Chromatography coupled to High-Resolution Mass Spectrometry (LC-HRMS) for polar metabolites of cells was carried out using a Q Exactive Plus.

### Liquid chromatography

A ZIC-pHILIC 150 × 2.1 mm (5 µm particle size) column (EMD Millipore) was employed on a Vanquish Horizon UHPLC system for compound separation at 40°C. The autosampler tray was held at 4°C. Mobile phase A is water with 20 mM Ammonium Carbonate, 0.1% Ammonium Hydroxide, pH 9.3, and mobile phase B is 100% Acetonitrile. The gradient is linear as follows: 0 min, 90% B; 22 min, 40% B; 24 min, 40% B; 24.1 min, 90% B; 30 min, 90% B. The follow rate was 0.15 ml/min. All solvents are LC-MS grade and purchased from Fisher Scientific.

### Mass spectrometry

The Q Exactive Plus MS (Thermo Scientific) is equipped with a heated electrospray ionization probe (HESI) and the relevant parameters include: heated capillary, 250°C; HESI probe, 350°C; sheath gas, 40; auxiliary gas, 15; sweep gas, 0; spray voltage, 3.0 kV. A full scan ranged from 55 to 825 (*m/z*) was used. The resolution was set at 70,000. The maximum injection time was 80 ms. Automated gain control (AGC) was targeted at 1 × 10^6^ ions. Maximum injection time was 20 msec. Raw data collected from LC-Q Exactive Plus MS was processed on Skyline (https://skyline.ms/project/home/software/Skyline/begin.view) using a five ppm mass tolerance and an input file of *m/z* and detected retention time of metabolites from an in-house library of chemical standards. The output file including detected *m/z* and relative intensities in different samples was obtained after data processing. Quantitation and statistics were calculated using Microsoft Excel, GraphPad Prism 8.1, and Rstudio 1.0.143.

### Clinical specimen analysis

Serum specimens were drawn from patients on Cycle 1 Day 15 at 0h, 0.5h, 1, 1.5h, 2h, 4h, 6h, 8h, 10h, 12h after administration of RGX-202 and were analyzed for RGX-202 using an analytically validated LC-MS/MS method at Syneos Health. Urine specimens were collected on Cycle 1 Day 15 at 0, 4 and 12 h after administration of RGX-202; average values are presented in the figure. Serum sample for creatine analysis was collected at 12h post-administration. Creatine levels were analyzed by Q2 solutions at contract research laboratories.

### Patient Details

In the RGX-202 Phase 1 a/b human study, adult patients of both sexes over the age of 18 years were enrolled. Data from 13 patients were analyzed. The study is currently accruing and ongoing. The protocol was approved by all site Institutional Review Boards and all patients signed informed consents before any screening procedures were obtained.

### Clinical Study Design

This Phase 1 a/b study is open-label, multi-center and single-arm, whose primary objective is to determine the maximum-tolerated dose, or maximum tested dose at which multiple dose-limiting toxicities (DLTs) are not observed, of RGX-202. Inclusion and exclusion criteria are stipulated, requiring all patients to have a pathologic confirmation of a locally advanced or metastatic solid tumor or lymphoma that has been deemed refractory to standard therapies. Patients may not have any other active malignancy that could confound the study endpoints. Patients must not have a history of pancreatitis, active Hepatitis B or C, or any illness/social situation in the opinion of the treating investigator that would limit compliance with the study requirements. Patients are not allowed to be treated with any other anti-neoplastic therapies while on study. Typical Phase 1 study parameters are required for performance status, hematologic and other organ function measurements. Patients are required to use acceptable contraceptive methods while on study and for a specified time period thereafter. Concomitant medications are restricted only if they pose a clinical risk of drug-drug interactions. Treatment for any condition with corticosteroids is not allowed unless at doses less than 10 mg daily prednisone equivalents. Patients were treated with RGX-202 at 600 mg, 1200 mg, 2400mg or 3600 mg twice/day continuous dosing, according to the patient cohort to which they were enrolled. Patients continued treatment with RGX-202 until treatment intolerance or progression of disease. The primary endpoint is incidence of DLTs, which are evaluated by the Rgenix sponsor medical monitor in collaboration with all treating clinical investigators. Secondary endpoints include pharmacokinetic measurements of RGX-202 and its metabolites in plasma and urine. Exploratory endpoints include measuring of serum and urine levels of creatine. Ultimately, efficacy endpoints will be obtained on a large sample size of patients in disease-specific expansion cohorts. Dose-limiting toxicities are defined as any of the following toxicities occurring during the first 4 weeks of treatment that are not clearly related to another cause (i.e., disease progression): any grade ≥3 non-hematologic AE, with the exceptions of Grade 3 nausea, vomiting, diarrhea, constipation, fever, fatigue, or skin rash in which there has been suboptimal prophylaxis and management that resolves to Grade % 2 within 72 hours; grade 4 thrombocytopenia, or Grade 3 thrombocytopenia with Grade > 1 bleeding or requirement for platelet transfusion; grade 4 neutropenia; grade ≥3 febrile neutropenia; grade ≥3 transaminase (AST/ALT) elevation; any toxicity resulting in > 25% held/skipped doses during the cycle; any other significant toxicity considered by the Investigator Sponsor’s medical representatives to be dose-limiting.

### Statistical analysis

Significance of tumor growth curve comparisons was carried out using two-sided t tests. Metastasis assays were analyzed using the non-parametric Mann-Whitney test. The Mantel-Cox log-rank test was used for statistical comparisons in survival analyses. Statistical analysis of creatine and RGX-202 AUC correlations in human patients and mice were carried out using simple linear regression with Prism 8. Statistical comparisons of cleaved Caspase-3 signal, in vitro and in vivo d3-creatine analysis, qPCR and IHC quantification were carried out using two-tailed t tests. T-test was used to compare metabolite abundance with Bonferroni’s multiple test correction where indicated. Throughout all figures: *p < 0.05, **p < 0.01, and ***p < 0.001, ****p < 0.0001. Significance was concluded at p < 0.05.

## Acknowledgments

We are grateful to members of our laboratories for providing insightful comments on past versions of this manuscript. We thank Eduardo Martinez for guidance on in vitro and in vivo creatine-d3 experiments and David Darst for intellectual and operational guidance in RGX-202 development. We thank Steve Wald for oversight of RGX-202-01 production and Michael Szarek for regulatory guidance. We thank Chelsea Bailey, Crystal Sun, Katharina Hodges and their teams at Crown Biosciences for performing the PDX trial. We thank Suresh Anaganti at AJES/L2P Research for conducting the in-life portion of the mouse creatine experiment. We thank Q2 solutions and Syneos Health for bioanalytical analysis.

## Funding

SKI 05160 General Fund (AB, KM)

NCI award CCSG P30 CA008748-53 (AB, KM)

Robertson Foundation Pre-Clinical Award (HT and NY)

Black Family Center for Research on Human Cancer Metastasis (NY)

The National Center for Advancing Translational Sciences UL1TR001866 (NY) Meyer Clinical Scholar Support (NY)

The Rockefeller University Clinical Scholar Endowment Fund (NY) Faculty Scholars grant from the Howard Hughes Medical Institute (SFT) The Black Family Foundation (SFT).

## Author contributions

Conceptualization: IK, MFT, SFT

Experimentation and analysis: IK, NY, CAA, HT, SS, ST, FG, JL, RB

Supervision: IK, CA, NY, FG, MFT, SFT

Design and supervision of the clinical trial: MFT, RW, SR, SLS, SFT

Clinical investigators: JCB, JS, MF, AJM, AEH, LSR, AC

Writing—original draft: IK, NY, CAA, SFT

Writing—review & editing: IK, ST

## Data and materials availability

All data associated with this study are available in the main text or the supplementary materials.

The data that support the findings of this study are available on request from the corresponding authors (IK, MFT, SFT).

## Supplementary Materials

**Fig. S1.**
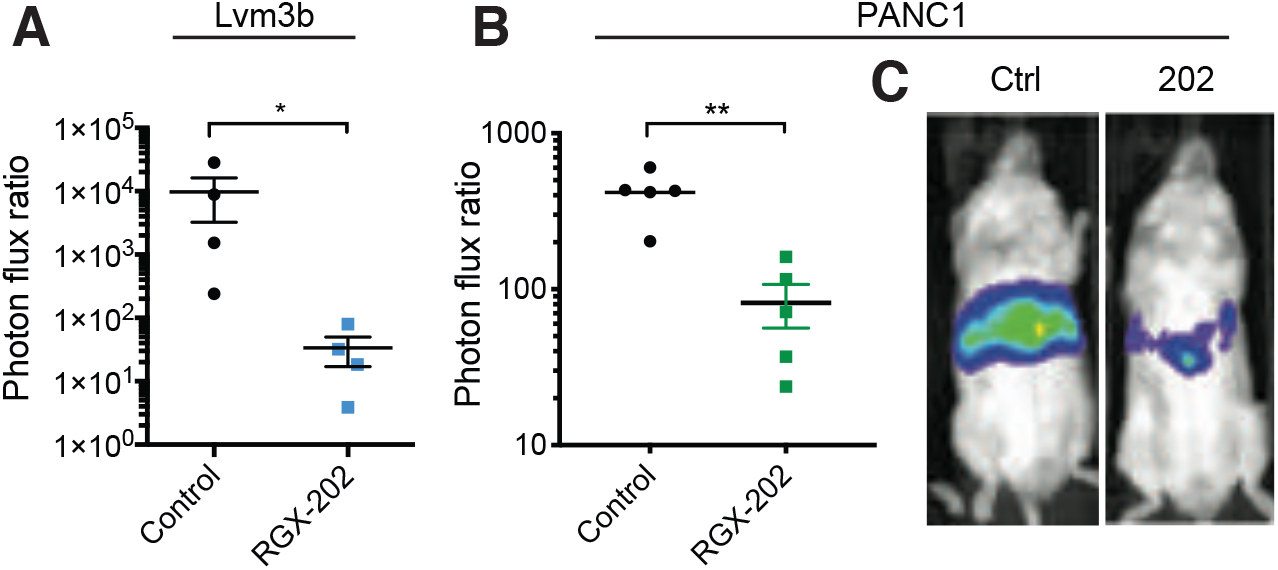
RGX-202 inhibits liver colonization of metastatic CRC and pancreatic cell lines. **(A)** Bioluminescence plot of liver colonization by 5×10^5^ human Lvm3b metastatic colon cancer cells after intra-splenic injections into NOD-SCID mice. Treatment with control or RGX-202-formulated diet started on day 1, for the duration of the experiment. Mice were imaged on day 22 after injection. Photon flux ratio is the ratio of bioluminescence signal at day 14 normalized to the signal on day 0; n = 4 per group, p = 0.0286. **(B, C)** Bioluminescence plot of liver colonization by 5×10^5^ human PANC1 pancreatic cells that were pre-treated with RGX-202 at 10 mM for 48h, after intra-splenic injections into NSG mice **(B)**. Photon flux ratio **(C)** is the ratio of bioluminescence signal at day 28 normalized to the signal on day 0; n = 5, p = 0.0079. All p values are based on Mann-Whitney test.

**Fig. S2.**
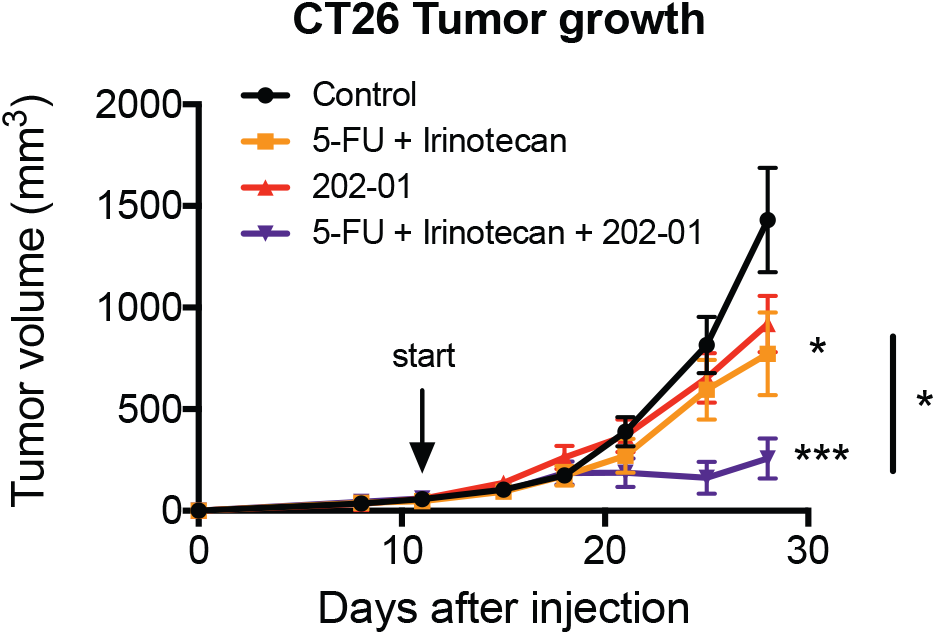
Anti-tumor efficacy of RGX-202 in combination with 5-FU and irinotecan. Tumor growth of 1.5×10^5^ CT26 cells subcutaneously injected into BALB/c mice. As soon as tumors reached an average of 50 mm^3^, mice were randomly distributed and received either a control diet, RGX-202-01-supplemented diet (∼500 mg/kg, p = 0.092), 5-FU (i.p. 30 mg/kg/week) and irinotecan (i.p. 15 mg/kg/week) (p = 0.064) or a combination of RGX-202-01, 5-FU and irinotecan (p = 0.0006); = 7-8 per group; All p values are based on two-sided t tests; *p = 0.02.

**Fig. S3.**
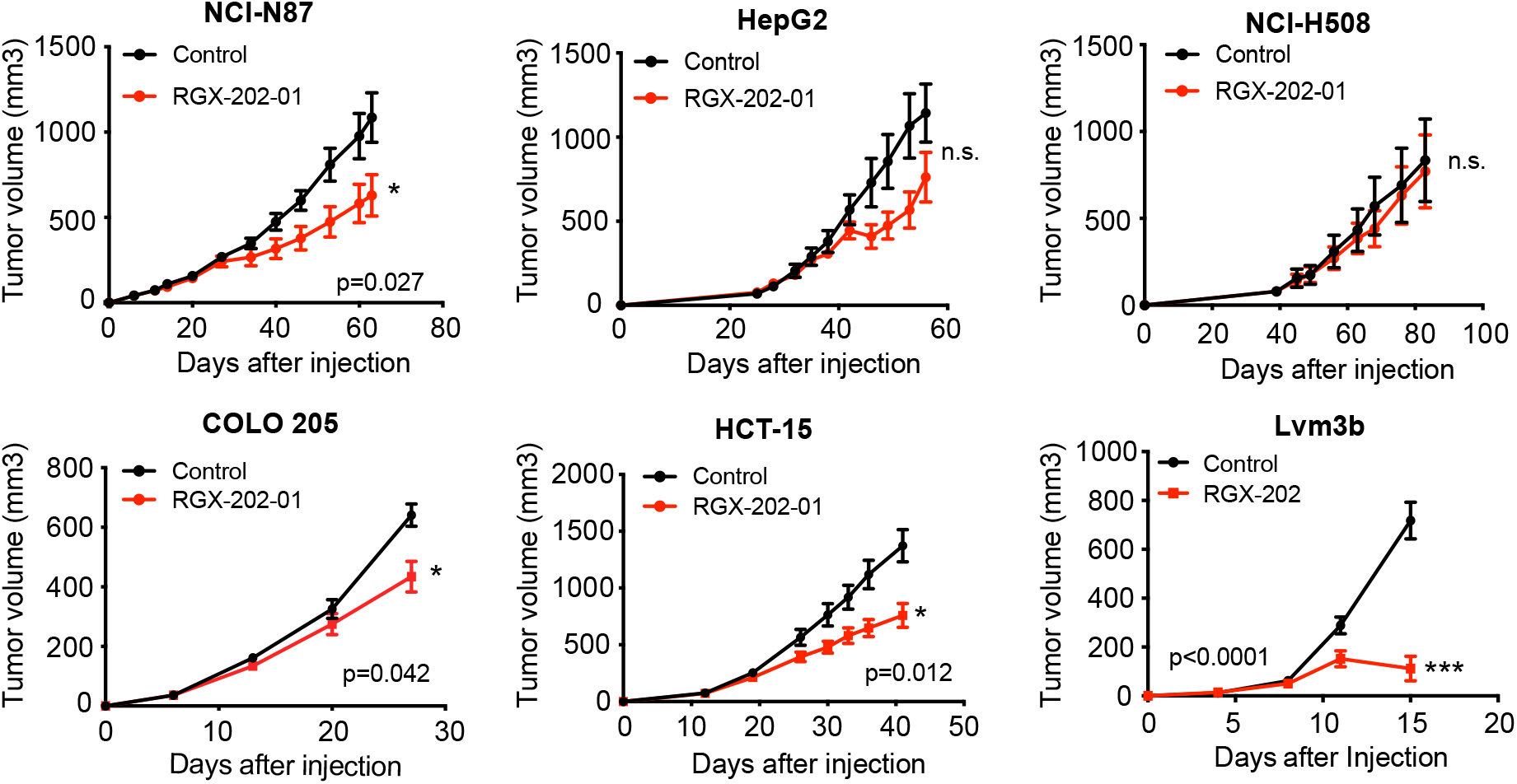
Xenograft tumor studies. Tumor growth of 4 x 10^6^ NCI-N87 (p = 0.027), 2 x 10^6^ HepG2, 2.5 x 10^6^ NCI-H508, 5 x 10^6^ Colo205 (p = 0.042), 2 x 10^6^ HCT-15 (p = 0.012) and 1 x 10^6^ Lvm3b (p<0.0001) cells subcutaneously injected into athymic nude mice. Treatment with a control or RGX-202/RGX-202-01-supplemented diet (∼800 mg/kg/day) started as soon as tumors reached 10-60 mm^3^; n = 8-10 per group. All p values are based on two-sided t tests; n.s. = not significant.

**Fig. S4.**
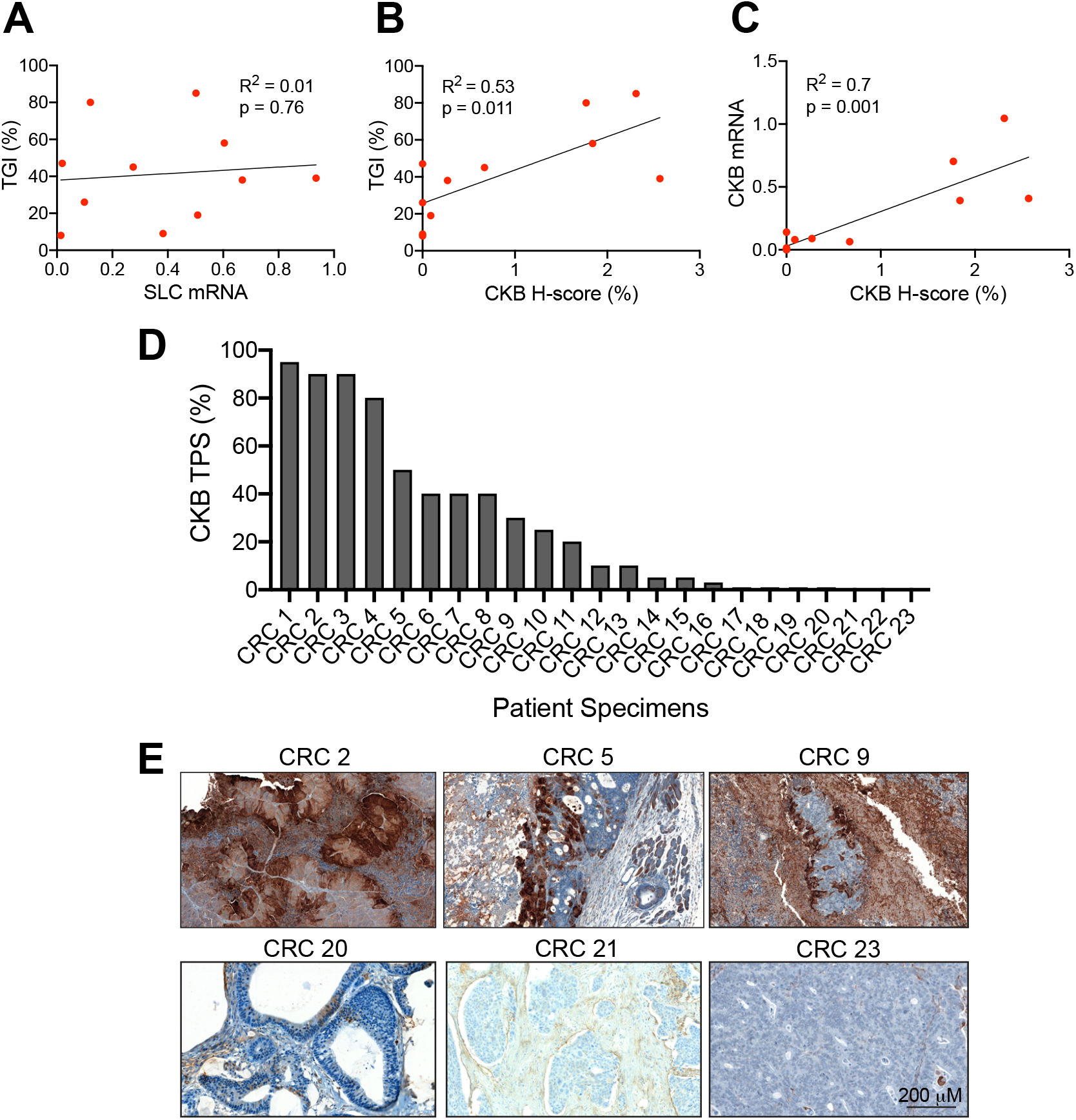
Linear regression analysis demonstrates anti-tumor efficacy of RGX-202 in CKB high expressing cell lines. **(A-C)** Linear regression analyses of the percentage of tumor growth inhibition (TGI) and the SLC6A8 mRNA **(A)**, TGI and CKB H-score **(B)** and CKB mRNA levels and CKB H-score **(C)** in the tumors of the xenograft models tested; n = 11. (**D, E)** CKB expression was assessed by IHC in 23 metastatic CRC specimens. Quantification of TPS is shown in **(D)** and representative images are shown in **(E)**.

**Table S1.** PDX models utilized in the PDX trial. Summary of PDX tumor growth data and KRAS status. TGI(21) = Tumor growth inhibition on Day 21 = ((C_21_-C_0_)/C_21_) – ((T_21_-T_0_)/T_0_). Change in tumor size relative to control.

| Cancer Type | ID | TGI(21) | Change in tumor size | KRAS | BRAF |
| --- | --- | --- | --- | --- | --- |
| CRC | CR5039 | -3.67 | 147% | WT | WT |
| CRC | CR3280 | -3.74 | 141% | p.G12R | D22N |
| CRC | CR5053 | -2.78 | 71% | p.A146V | WT |
| CRC | CR2554 | -1.03 | 44% | p.G12V | WT |
| CRC | CR5032 | -1.91 | 41% | p.G13D | WT |
| CRC | CR6251 | -2.79 | 29% | WT | WT |
| CRC | CR5066 | -0.61 | 24% | p.G12C | WT |
| CRC | CR5075 | -1.23 | 17% | WT | V600E |
| CRC | CR5033 | -0.47 | 14% | WT | WT |
| CRC | CR5080 | 0.50 | 6% | p.G12D | WT |
| CRC | CR5081 | -0.15 | 1% | p.G12D | WT |
| CRC | CR1287 | 0.04 | 0% | p.G12D | WT |
| CRC | CR5059 | -0.18 | -2% | WT | V600E |
| CRC | CR3151 | 0.23 | -2% | p.G12A | WT |
| CRC | CR2491 | 0.26 | -4% | p.G12V | WT |
| CRC | CR5052 | 0.11 | -5% | WT | V600E |
| CRC | CR5046 | 0.43 | -8% | p.G12R | WT |
| CRC | CR6460 | 2.26 | -14% | p.Q61L | WT |
| CRC | CR5082 | 0.35 | -16% | WT | V600E |
| CRC | CR6863 | 0.55 | -19% | WT | V600E |
| CRC | CR5043 | 0.57 | -26% | p.G13D | WT |
| CRC | CR3085 | 3.28 | -27% | WT | WT |
| CRC | CR5058 | 1.81 | -32% | WT | WT |
| CRC | CR5044 | 1.69 | -32% | WT | WT |
| CRC | CR5048 | 1.00 | -33% | p.G12C | WT |
| CRC | CR5038 | -0.07 | -34% | p.Q61H | WT |
| CRC | CR5101 | 1.10 | -36% | WT | WT |
| CRC | CR2526 | 1.01 | -42% | WT | WT |
| CRC | CR5040 | 6.75 | -42% | p.G12V | WT |
| CRC | CR2179 | 3.34 | -45% | WT | V600E |
| CRC | CR6249 | 3.08 | -45% | WT | WT |
| CRC | CR11372 | 1.79 | -47% | WT | WT |
| CRC | CR6256 | 3.82 | -48% | p.G12C | WT |
| CRC | CR1795 | 9.37 | -48% | WT | WT |
| CRC | CR5026 | 2.94 | -49% | WT | V600E |
| CRC | CR3079 | 3.01 | -49% | p.G12D | WT |
| CRC | CR3099 | 4.05 | -49% | p.G12D | WT |
| CRC | CR2533 | 4.18 | -51% | WT | WT |
| CRC | CR5030 | 4.43 | -51% | WT | WT |
| CRC | CR3448 | 5.94 | -56% | p.G12D | WT |

**Table S2.**
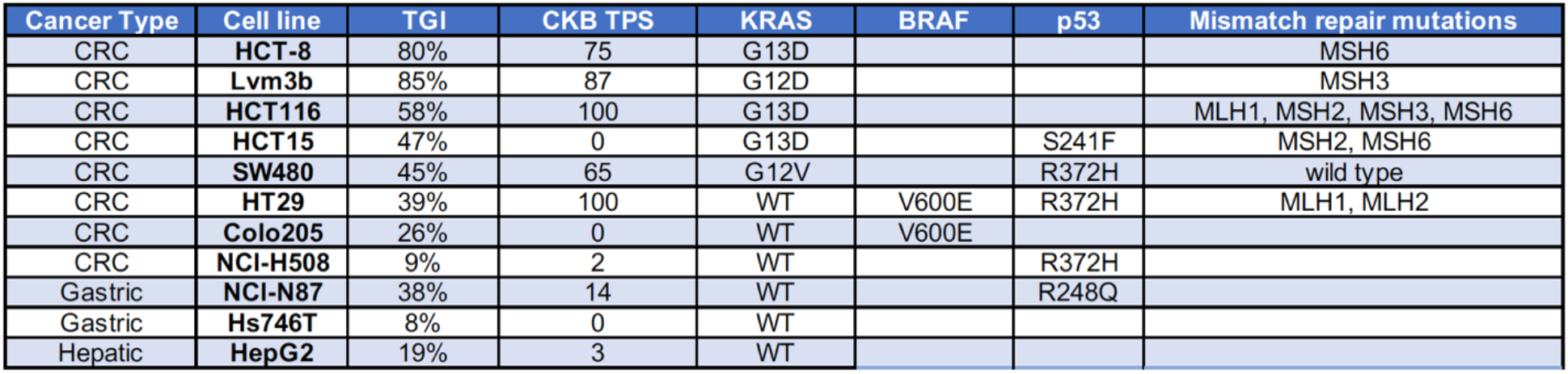
Information on cell lines. Anti-tumor efficacy and mutational status of the cell lines used in the xenograft studies. TGI = Tumor growth inhibition.

